# Rational cell culture optimization enhances experimental reproducibility in cancer cells

**DOI:** 10.1101/167148

**Authors:** Marina Wright Muelas, Fernando Ortega, Rainer Breitling, Claus Bendtsen, Hans V. Westerhoff

## Abstract

Optimization of experimental conditions is critical in ensuring robust experimental reproducibility. Through detailed metabolomic analysis we found that cell culture conditions significantly impacted on glutaminase (GLS1) sensitivity resulting in variable sensitivity and irreproducibility in data.

Baseline metabolite profiling highlighted that untreated cells underwent significant changes in metabolic status. Both the extracellular levels of glutamine and lactate and the intracellular levels of multiple metabolites changed drastically during the assay. We show that these changes compromise the robustness of the assay and make it difficult to reproduce.

We then devised ‘metabolically rationalized standard’ assay conditions, in which glutaminase-1 inhibition reduced glutamine metabolism differently in both cell lines assayed, and decreased the proliferation of one of them. The adoption of optimized conditions such as the ones described here should lead to an improvement in reproducibility and help eliminate false negatives as well as false positives in these assays.

## 1 Introduction

Reproducibility has increasingly become a topic of concern in biomedical research^1,2^. Scientists acknowledge that they fail to reproduce even their own experiments, let alone those of their colleagues around the globe^3^. When testing a potential anticancer drug, a novel and potent allosteric inhibitor specific for the glutaminase-1 enzyme (EC 3.5.1.2), we initially experienced a similar irreproducibility. Our focus on metabolomics led us to experiments that then produced an explanation for the lack of reproducibility, and employed a more comprehensive assay development approach which we believe can be of benefit for the scientific community.

One of the initial steps in the development of therapeutic agents for cancer involves testing these agents *in vitro* using human cancer cell lines as experimental models^4,5^. Using primary cell lines in culture, the effects of compounds or perturbations on cell proliferation, DNA replication or cell death is generally investigated over a period of time. These types of read-out are highly dependent on cell physiology and as such these assays need to fulfill a number of conflicting conditions. On the one hand, cells need to be kept in culture long enough to attain a steady state and for the effects of treatments to be observed. On the other hand, they should not be kept there too long because of the gradual accumulation of waste products that can be inhibitory or toxic to cells, such as lactate and ammonia^6^. The concentration of nutrients will fall over time, pH will change, and as cells grow and divide, space may become limiting. As cell density increases, effects of paracrine signaling become more pronounced and as cells reach confluence, contact inhibition may suppress proliferation. Although cancer cells are able to proliferate for some time after reaching confluence by then accumulating on top of one another, this crowding still limits individual cells’ access to nutrients and growth factors^7^, eventually resulting in cell cycle arrest and apoptosis, but long before then, in shifts in cell metabolism. Cell viability assays are affected by the metabolic state of the cells and therefore any shift in metabolic states during the assay, and particularly different shifts between sensitive and resistant cell lines, would confound the outcome of such assays.

Recently, Haibe-Kains *et al.* highlighted multiple inconsistencies between two large-scale pharmacogenomic studies, the Cancer Genome Project (CGP^8^) and the Cancer Cell Line Encyclopedia (CCLE^9^), *viz.* the sensitivity profiles of common cell lines and drugs^10^. It has been suggested that differences in the cell culture conditions were amongst the reasons for these discrepancies^11^ and that consistency should be achievable with appropriate laboratory and analysis protocols^12^. For example, for each cell line the CGP study determined the seeding density that ensured that each was still in the growth phase at the end of the assay (∼70% confluence), whilst the seeding density was not reported for the CCLE study^8^. In addition, for adherent cells, the test compound was added ‘around 12–24 hours’ after seeding cells and studied over a further 72–84 hours in the CCLE study, whereas in the CGP study this was added 1 day after seeding and assayed 72 hours after treatment. This lack of standardized and well-described culture conditions is common to most literature in this field (see refs. ^13-19^ for examples). Living cells are complex; they adjust to altering environments, by quick metabolic or somewhat slower gene-expression regulation and this may readily change the extent to which any target limits cell physiology and survival. This can make results of drug targeting studies irreproducible unless the relevant environmental conditions are well controlled at the appropriate time scale. Both academia and the pharmaceutical industry recognize the necessity of much more thorough standardization to improve reproducibility ^20^.

The metabolic performance of the cell lines during drug targeting assays is not assessed routinely, or at least not reported. Metabolic changes could have strong implications for therapeutic targets in, or affected by, intermediary metabolism. Metabolic enzymes involved in cellular proliferation and growth have been identified as altered in cancers, either through the expression of cancer-specific isoforms, through mutations, or through altered expression levels^21^. And it is precisely these targets that are witnessing revived interest of late^22,23^: these altered metabolic pathways are now being targeted directly, used to enhance the efficacy of existing therapeutic agents or to overcome resistance to current treatment strategies for cancer. In addition, anti-cancer drugs that do not target metabolism itself are often assayed in survival based assays. If metabolism is so involved in cell survival, its variability and during survival based assays could therefore be a prime cause of irreproducibility of the outcome of the many experimental assays. We thus investigated whether variability in cellular metabolic status is linked with different phenotypic responses.

Here we show how culture conditions widely used to investigate the effects of an inhibitor of the glutaminase-1 enzyme on cell proliferation and metabolism, result in drastic and rapid changes in the metabolic state of the cells, compromising the robustness and reproducibility of the results. We then present the pipeline we engage in such cases in order to identify these changes and to optimize culture conditions accordingly. The reward is a robust study of the effects of a potentially important anti-proliferative agent.

## 2 Results

To investigate the effect of an inhibitor (GLS1i) of the glutaminase-1 enzyme (GLS1, EC 3.5.1.2), on cell metabolism and proliferation, we started by employing culture conditions that are widely used in the scientific literature for proliferation assays and should enable the application of metabolomics^13-19^. We seeded cells at a density of 8 × 10^5^ cells/well in 1 mL of culture media 24 hours prior to commencing the experiment at a time point denoted as time 0 by adding 1.0 μM of a GLS1 inhibitor (see Materials and Methods section 1.2). The effect of this inhibitor on cell survival was determined 48-hours later. We used two cell lines, A549 and H358, that are dependent on glutamine for proliferation^24^, but differ in sensitivity to a novel and potent inhibitor of GLS1 activity developed jointly by AstraZeneca and Cancer Research Technology: proliferation of A549 cells is inhibited by this ‘GLS1i’, whereas proliferation of H358 cells is insensitive to GLS1 inhibition (Figure S1).

### 2.1 The problem: the inhibitor does not seem to work

We had expected that treatment with a GLS1 inhibitor would lead to a reduced consumption of glutamine, a reduced production of glutamate, an increased intracellular concentration of glutamine, a reduced intracellular concentration of glutamate and reduced intracellular concentrations of all TCA cycle intermediates, particularly in the GLS1i sensitive A549 cell lines. The initially observed effects of GLS1i treatment were very different to what we expected (Table 1): GLS1i treatment did not affect cell numbers in either cell line when compared to control treatment (Figure S2a). Equally unexpectedly, the amount of glutamine consumed was reduced to a much greater extent in the resistant cell line than in the sensitive cell line (Figure 1a). Intracellular glutamine concentrations were raised in treated conditions in both cell lines (Figure 1b), particularly in the resistant H358 cell lines compared to controls, in agreement with our expectations. However, intracellular glutamate concentrations were reduced in the GLS1i resistant H358 cell lines only (Figure 1c). The abundance of TCA cycle intermediates, such as alpha-ketoglutarate (α-KG), citrate and fumarate, was unaffected by treatment of A549 cells with GLS1i. Only α-KG was reduced in H358 cells (Figure S3).

**Table 1.**
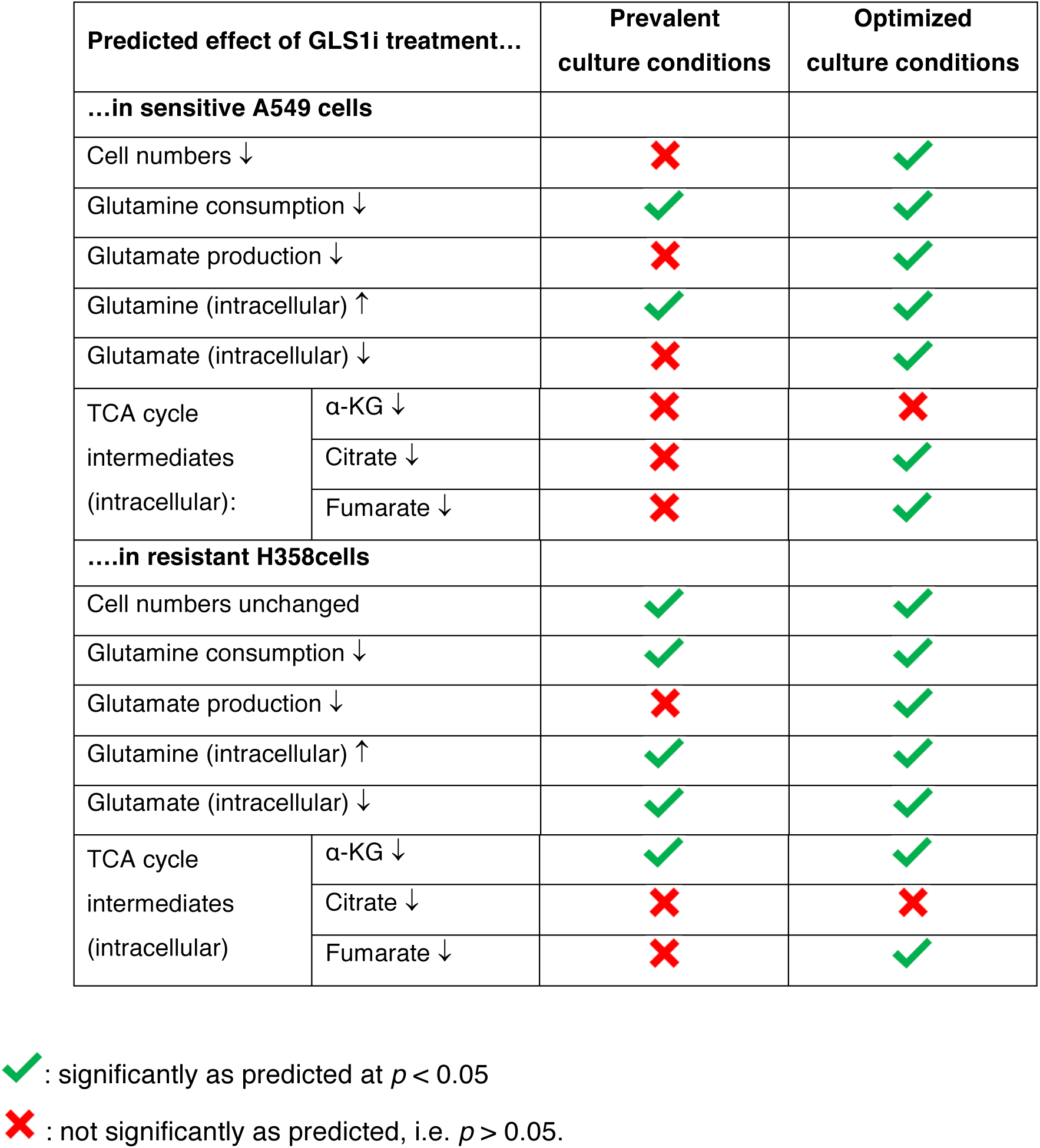
Predicted versus observed effects of treatment with the GLS1i inhibitor using the prevalent culture conditions compared with the optimized culture conditions devised in this study.

**Figure 1:**
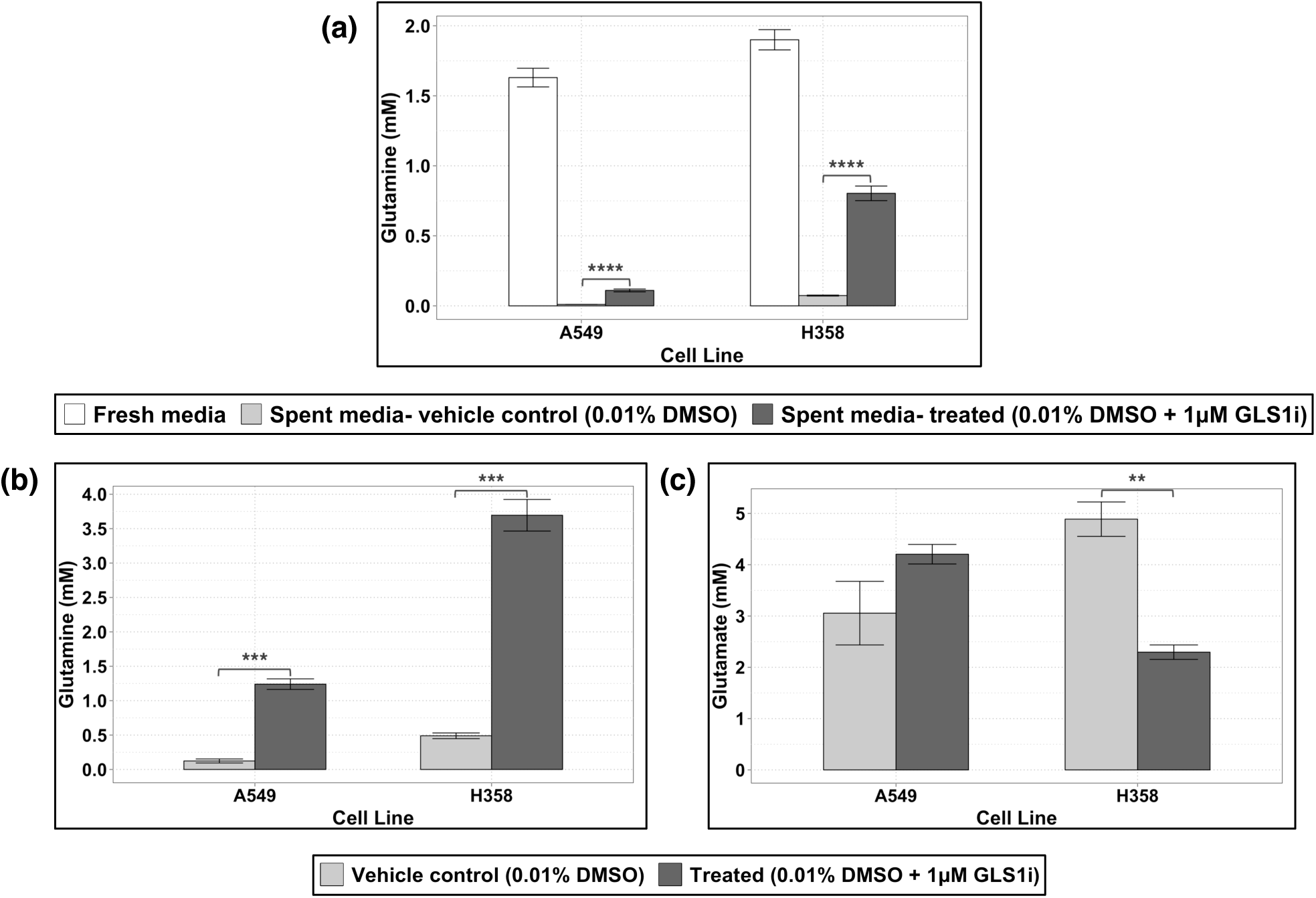
Glutamine and glutamate concentrations 48 hours after treatment with 0.01% DMSO ± 1 μM GLS1i using a prevalent assay method. A549 and H358 are known as sensitive and resistant cell lines, respectively, vis-à-vis glutaminase 1 inhibitors. Concentrations of **(a)** extracellular glutamine **(b)** intracellular glutamine and **(c)** intracellular glutamate measured by LC-UV after 24 h of treatment with 0.010 % DMSO ± 1.0 μM (final concentrations) GLS1i. For this single experiment, measurements were performed in triplicate for control and treated conditions. Cells had been seeded at a density of 8x10^5^ cells/well in 1 ml of culture media 24 h prior to commencing the experiment. Shown are the mean ± SEM for the 3 technical replicates per cell line and treatment condition. Unadjusted p-values of the differences between control and treated samples obtained using a two-tailed Student’s t-test are denoted with asterisks: **: p ≤ 0.01; ***: p ≤ 0.001; ****: p ≤ 0.0001.

A first clue on what could be responsible for the lack of effect of the metabolic inhibitor on the cell proliferation and the paradoxical effects on metabolism, was the extracellular concentration of glutamine at the end of this assay: this was very close to undetectable levels, suggesting that during the assay cells had been subject to a concentration of glutamine varying between 2mM and 0 mM. In the absence of glutamine, an inhibitor of glutaminase should perhaps not be expected to have any effect; either directly or due to metabolic rewiring.

### 2.2 Explanation: The cellular environment is uncontrolled

To understand why GLS1i treatment in the above assay failed to show any significant effects on proliferation or metabolism by A549 and H358 cells we examined the changes in cell numbers and intracellular and extracellular metabolites with enhanced time resolution (Figure 2 and Figure S4).

**Figure 2:**
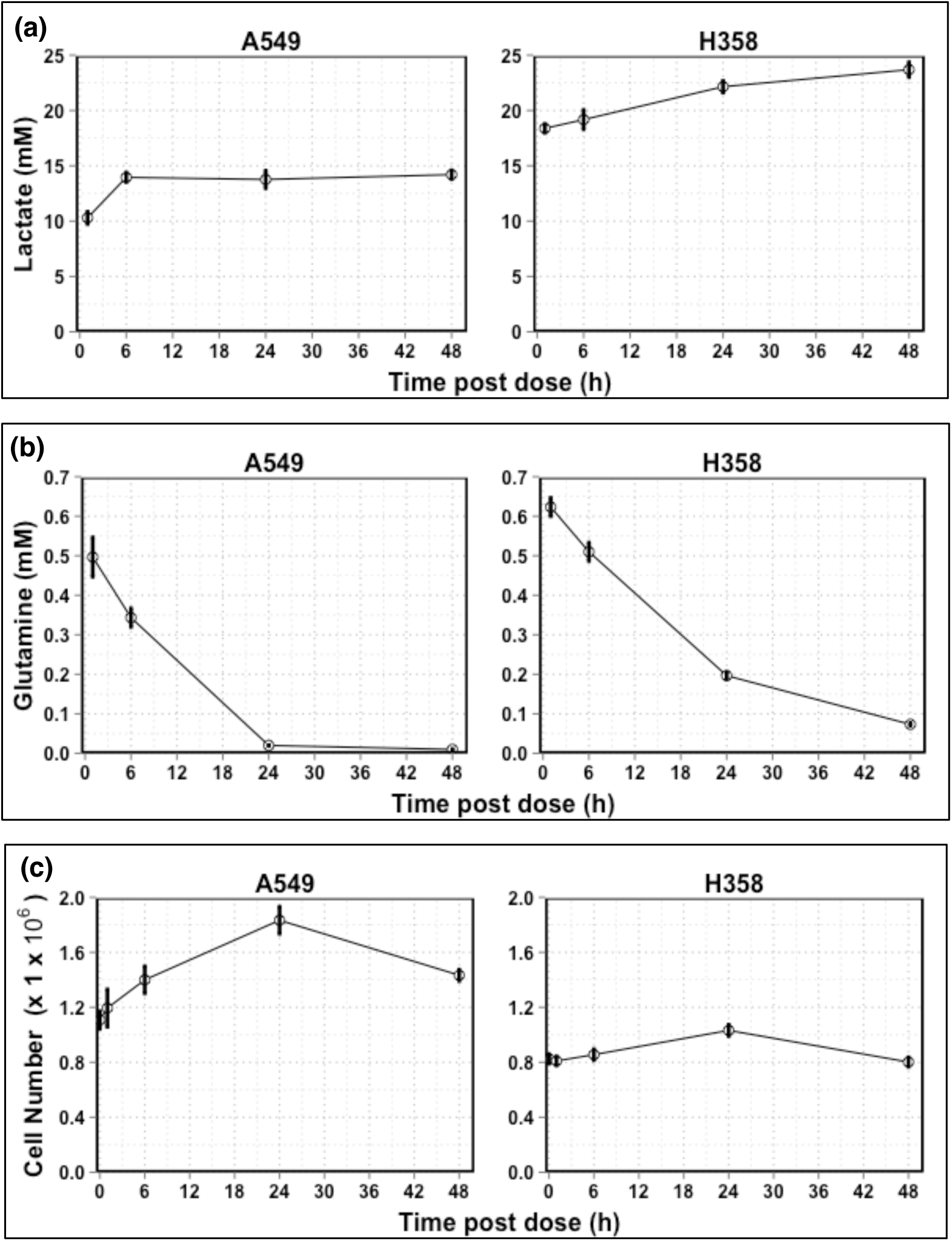
Changes with time after seeding of the state of a cell culture in a traditional assay in vehicle control (0.01% DMSO) and treatment conditions. **(a)** Lactate (measured by LC-MS) and **(b)** glutamine (measured by LC-UV) in spent medium. **(c)** Number of live cells per well as measured using the Trypan blue exclusion technique using a Countess automated cell counter (Thermo Scientific, Loughborough, UK). Zero time corresponds to 24 hours after seeding of cells into a medium containing 10 mM of glucose, 2 mM of glutamine, in addition to dialyzed fetal calf serum, vitamins and both essential and non-essential amino acids at concentrations well below 1mM except for arginine (0.95 mM) and glutamine (2 mM), as shown in Table M1 in Materials and Methods. The cell lines were: A549 (left) and H358 (right). For this single experiment, measurements were performed in triplicate. Shown are the mean ± SEM for the 3 technical replicates per cell line in control conditions.

Our results suggest that, throughout the course of the assay, the cells’ environment in control conditions was changing in ways that would be expected to interfere with the cells’ internal state^25^. Firstly, the concentration of lactate in spent media 1 hour into the assay was already above 10 mM in both cell lines (Figure 2a). H358 cells had already secreted nearly 20 mM of lactate by this point, suggesting that most of the glucose available in culture media had already been consumed. In H358 cells this increase in lactate continued over the time points sampled, but in A549 cells the concentration of lactate reached its maximum level of around 14 mM 6 hours post dosing. Secondly, the concentration of glutamine in spent media was already reduced by ≥ 70% 1 hour into the assay in both cell lines and undetectable by 24 hours and 48 hours post dose in A549 cells and H358 cells, respectively (Figure 2b).

The fluctuating environment that the cells were exposed to in this assay likely contributed to the changes in the specific growth rate of these cells: a small increase in cell numbers was observed over the first 24 hours post dosing (Figure 2c) but was much slower than the expected growth kinetics of these two cell lines^26^. Moreover, between 24 and 48 hours post dosing, the number of cells in control conditions seemingly *decreased*. This could be due to the depletion of glutamine, glucose or other essential substrates not measured, to the increases in the concentration of lactate, to the resulting decrease in pH, or to contact inhibition of the cells. Deprivation of nutrients and growth factors has been shown to lead to cell cycle arrest and subsequently cell death in NSCLC cell lines indicating that this is a possible explanation for the changes seen in cell numbers in this type of assay^27-29^. The drastic reductions in glutamine could influence normal cell metabolism and physiology, as a result of forcing cells to switch to alternative fuel sources and to deal with the problem of ammonium toxicity^6^. Indeed, the intracellular concentration of glutamine fell drastically throughout the assay, as did the abundance of other metabolites, albeit to a smaller extent (Figure S4). The high concentration of secreted lactate in spent media is likely to be accompanied by drastic acidification of culture media and cellular damage^30,31^; the culture media had a buffering capacity of around 20 mM/pH unit, whilst some 20 mM of lactic acid may have been produced, a large proportion of which was likely derived from glucose.

Our results suggest that these commonly used assay conditions are unsuitable for comparing inhibitors of molecular targets with each other.

### 2.3 Assay optimization: Reducing the seeding density and increasing culture volume stabilizes cellular state

Increasing the volume of culture media alone from 1 to 3 mL was not sufficient to avoid these problems of variations in metabolic state (Figure S5): Whilst this reduced the magnitude of changes in the extracellular concentrations of glucose and glutamine, these key nutrients were still close to depletion 72 hours after seeding. pH changes remained within acceptable ranges in A549 cells, but not in H358 cells where pH changed by > 1 pH unit. A slight improvement in the proliferation of these two cell lines was observed but this was still much slower than expected. Confluence was reached early into the assay (24–36 hours after seeding) when the cell lines were seeded at a density of 8 × 10^5^ cells/well (Figure 3, upper purple line). This, together with the drastic reductions in nutrient concentrations through the assay, may account for the reduced rate of proliferation observed in our previous assays as a result of the induction of cell cycle arrest and apoptosis^7,27-29,32^.

**Figure 3:**
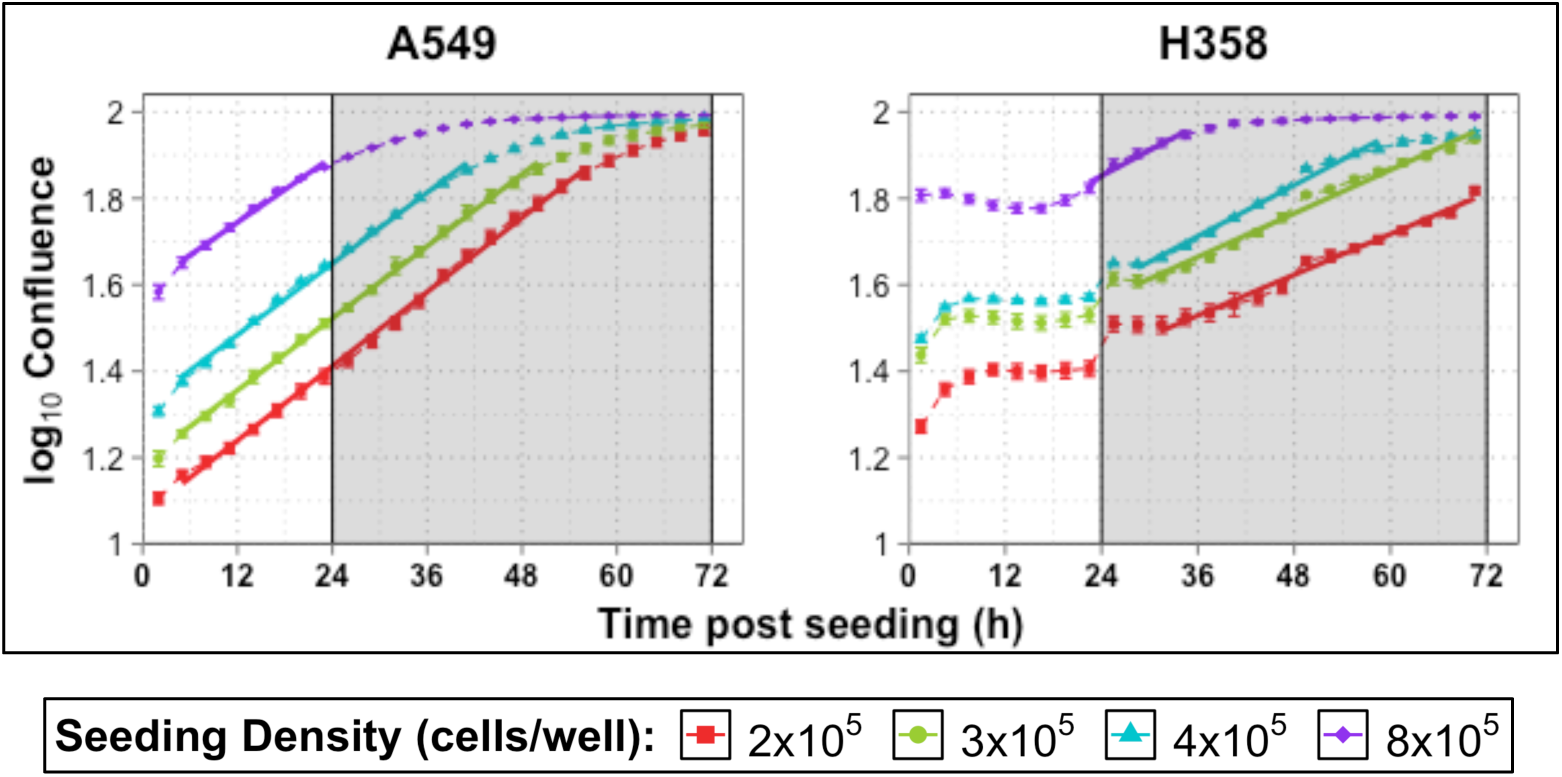
Confluence of A549 and H358 cells over 72 hours after seeding at different initial densities when the volume of culture media was increased to 3 mL. Shown are the changes in log_10_ confluence over time measured by live content cell imaging Incucyte HD system (Essen Bioscience) when A549 and H358 cells were seeded at a density of 8 × 10^5^, 4 × 10^5^, 3 × 10^5^ and 2 × 10^5^ cells/well. For this single experiment, measurements were performed in triplicate for control and treated conditions. Shown are the mean ± SEM for the 3 technical replicates per cell line and treatment condition. Shaded area denotes the assay window in a prevalent assay where samples would be taken over 48 hours from the time of dosing (24 hours after seeding). Solid lines are a fitted linear model for the exponential growth phase of cells.

Indeed, we observed steadier metabolic conditions and cell proliferation when the initial seeding density of cells was reduced *and* the volume of culture media increased from 1 to 3 mL (Figure 3 and Figure S5). The period of time during which cells were able to grow exponentially was also increased (Figure 3). Ensuring that confluence remained below ∼ 80 % throughout the assay window (24–72 hours post seeding), or that this level of confluence was reached as late as possible in the assay, required a significant reduction in the initial seeding density of cells, and this was cell-line specific. The time required to recover from reseeding also differed between cell lines and was affected by the initial seeding density. This initial lag phase was very short in duration for A549 cells (< 6 hours) compared to the approximately 24 hours required by H358 cells (Figure 3), which extended beyond 24 hours when H358 cells were seeded at 2 × 10^5^ cells/well. These differences in growth kinetics could well compromise inhibitor assays.

Lowering the initial seeding density of cells also reduced the magnitude of changes in the concentrations of key nutrients such as glucose and glutamine (Figure S5b and c), and in pH (Figure S5a) throughout the assay window (24–72 hours post seeding). In the case of the H358 cell line, using these conditions, assays beyond 48 hours after seeding may not be suitable since these cells displayed a high rate of glucose consumption (Figure S5b) and the corresponding lactate secretion would lead to significant reductions in pH (Figure S5a). When H358 cells were seeded at 3 x 10^5^ cells/well, the concentration of glucose reached limiting levels (∼2 mM) 72 hours after seeding, which would constitute 48 hours post dosing in an assay where treatment was applied 24 hours after seeding (Figure S5b).

We conclude that, in order to ensure that (1) cells are in exponential growth from 24 hours after seeding, (2) confluence is reached as late as possible, and (3) changes in glucose, glutamine and pH are kept to a minimum, the volume of culture media should be increased up to 3 mL and seeding density reduced according to individual cell line growth kinetics. In our case, seeding A549 cells at a density of 2 × 10^5^ cells/well or less, and H358 cells at around 3 × 10^5^ cells/well in 3 mL of culture media, fulfills these criteria.

### 2.4 Optimized in vitro culture conditions enable successful hypothesis validation and discovery

To validate the expected improvement in assay performance we then seeded A549 and H358 cells at a density of 1.5 × 10^5^ and 3 × 10^5^ cells/well respectively in 3 mL of media in a 6 well plate format. Cells were growing exponentially at rates comparable to those reported in the literature^26^ (Figure S6) throughout the assay in control conditions. From plates prepared in parallel, the levels of various metabolites in cell and spent media extracts as well as cell numbers were measured for 24 hours after treatment with 1 μM of the GLS1 inhibitor. In agreement with our expectations (Table 1), treatment with the GLS1 inhibitor over 24 hours led to a reduction in cell numbers of around 20% in A549 cells but not in H358 cell lines (Figure 4a). Throughout the assay, the changes in the cells’ environment were now minimal in both cell lines regardless of treatment conditions (Figure 4b-f): the concentrations of glucose and glutamine were reduced by less than 50% over the assay and the lactate secreted caused a pH drop < 1 unit under these improved assay conditions. The amount of glutamine consumed appeared reduced in both cell lines by treatment with the GLS1 inhibitor although these changes were small and only statistically significant in A549 cells: the achieved stability of culture conditions had the consequence that differences in cellular metabolism were no longer strongly reflected in the changes of the exometabolome, such that assay conditions were now under control and steady.

**Figure 4:**
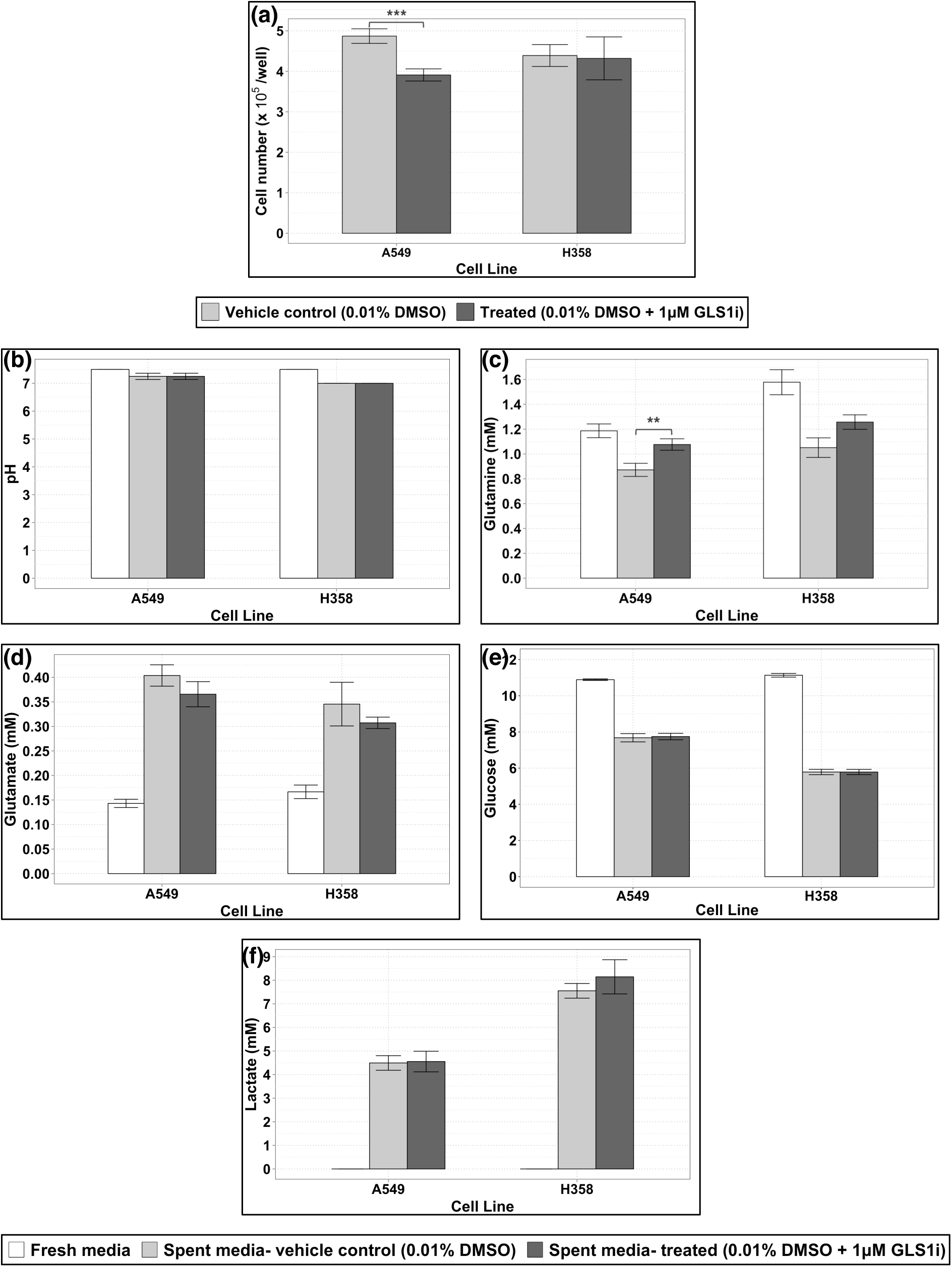
Live cell numbers, concentrations of various metabolites, and pH in fresh and spent media extracts 24 hours after treatment with 0.010 % DMSO ± 1.0 μM GLS1i using the improved culture conditions devised here. **(a)** Live cell numbers as measured by automated microscopy following Hoechst staining and fixation (See Materials and Methods 1.6). **(b)** pH of fresh and spent media samples measured using MColorpHast indicator strips. Concentration of **(c)** glutamine, **(d)** glutamate, **(e)** glucose and **(f)** lactate in fresh and spent media samples after treatment with or without 1.0 μM GLS1i measured by LC-UV (glutamine and glutamate), Accu-Chek Aviva Blood Glucose Meter System (glucose) and LC-MS (lactate). For each experiment, measurements were performed in triplicate for control and treated conditions. Shown are the mean ± SEM of 3 (A549 cell line; 9 data points per condition) or 2 (H358 cell line; 6 data points per condition) independent experiments. Note that glutamine concentrations in fresh media used for A549 and H358 cells fell by an average of ∼27 % and ∼5 % respectively over the duration of the assay. Unadjusted p-values of the differences between control and treated conditions obtained using a two-tailed Student’s t-test are denoted with asterisks: **: p ≤ 0.01; ***: p ≤ 0.001.

We therefore assessed intracellular metabolism to investigate whether, under the optimized conditions, the predicted effects of GLS1i on intracellular metabolites were observed that had not been observed under the previous unstable conditions (Table 1). Our results confirm that the glutaminase inhibitor engaged with the intended target: large reductions (*p* < 0.01) in glutamate were observed in both cell lines (Figure 5a). Only minor increases in the concentration of glutamine were seen, probably as a result of rapid equilibration with the external medium via the glutamine transporter (Figure 5b). The intracellular abundance of TCA cycle intermediates was also affected by GLS1i treatment in both cell lines (Figure 5c-e).

**Figure 5:**
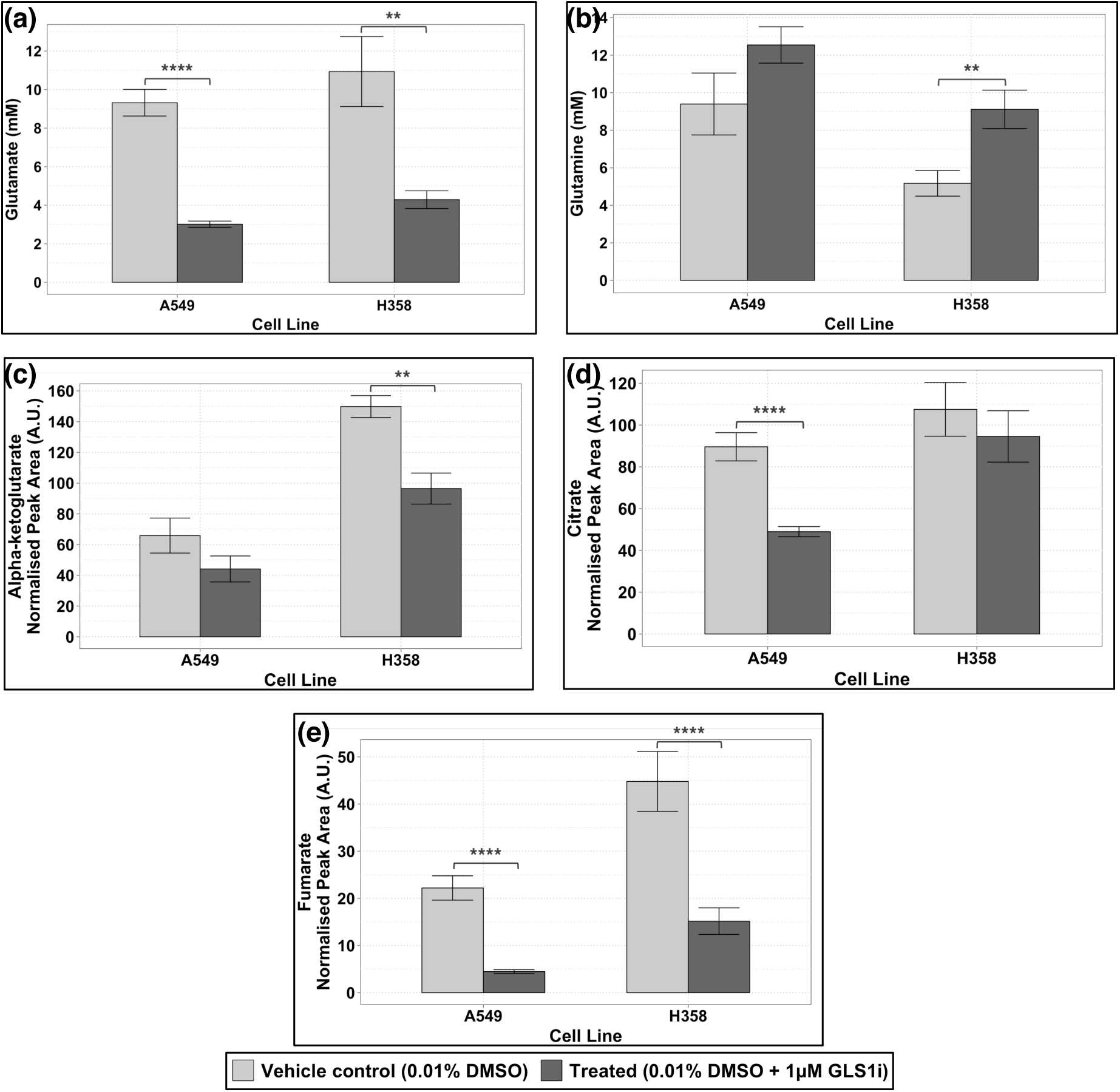
Levels of various intracellular metabolites 24 hours after treatment with 0.01% DMSO ± 1 μM GLS1i using the improved culture conditions devised here. A549 and H358 are known as sensitive and resistant cell lines, respectively. Concentration of **(a)** glutamate and **(b)** glutamine measured by LC-UV, and relative abundance of the TCA cycle intermediates (measured by LC-MS) **(c)** alpha-ketoglutarate, **(d)** citrate and **(e)** fumarate in cells after treatment with or without 1μM GLS1i. For each experiment, measurements were performed in triplicate for control and treated conditions. Shown are the mean ± SEM of 3 (A549 cell line; 9 data points per condition) or 2 (H358 cell line; 6 data points per condition) independent experiments. Unadjusted p-values of the differences between control and treated conditions obtained using a two-tailed Student’s t-test, the results of which are denoted with asterisks: **: p ≤ 0.01; ***: p ≤ 0.001; ****: p ≤ 0.0001.

Our results confirm that the optimized culture conditions devised here provide a robust and stable environment in which to reproducibly assay the effects of a GLS1 inhibitor on cell metabolism and proliferation.

## 3 Discussion

The assay with which we started this study failed to demonstrate any consistent effects of glutaminase inhibition on either glutamine metabolism or proliferation in these two cancer cell lines: addition of the inhibitor to cells (A549) known to be sensitive to the inhibitor, had no apparent effect on their proliferation (Figure S2a). Conversely, the glutamine metabolism by cells (H358) that are *in*sensitive to the same inhibitor was reduced to a much greater extent than that by sensitive (A549) cells (Figure 1). We then demonstrated that these inconsistencies were artifacts, for one, because most glutamine had been depleted in the pre-incubation period (Figure 1b and 2b) leaving too little glutamine for effects of the inhibitor to become statistically noteworthy.

With our improved assay conditions we were able to show that the glutaminase inhibitor (GLS1i) does have an effect on glutamine metabolism of both cell lines, but that only the proliferation of the A549 cells is reduced (Figure 4). GLS1 inhibition was also apparent from the changes in levels of intracellular metabolites in both cell lines, but with distinct differences between sensitive (A549) and resistant (H358) cell lines (Figure 5).

Our findings highlight the importance of *in vitro* assay optimization for the assessment of the potential of metabolic, and probably also other, inhibitors as anti-cancer drugs that impact on cellular metabolism. Variability in the metabolic state during the assay may well create false positives and false negatives because intermediary and energy metabolism is full of pleiotropic implications. Importantly perhaps, the implications of our findings are unlikely to be limited to studies of metabolic inhibitors. Other inhibitors, such as those of cell signaling or transcription require even longer cell incubations, and may therefore be compromised even more by changes in the levels of metabolites such as ATP, NADH, acetyl-CoA and glutamate that cross-talk widely. Even though metabolism may not be the drug target in these cases, its perturbation due to inappropriate culture conditions, might produce a false response. And since the drug may well affect metabolism indirectly, the impact of metabolic status could be overlooked in both control and treated conditions.

Perhaps even more so than this, our results should warn against the straightforward implementation of historically-fixed sets of conditions for drug assays in cell lines. Living cells are complex enough to engage in all sorts of metabolic changes, these changes may well differ between individual cell lines, and the metabolome is sensitive to such changes well before the metabolic fluxes produced by the cells are^33^. We therefore advocate that reports on drug assays are accompanied by a thorough description of the experimental procedure used as well as by a metabolic characterization of the cells during the assay, such as in the workflow we demonstrated here. After all, such characterization has become possible over recent years.

Indeed, reports in the literature regarding the characteristics of cell lines under basal and perturbed conditions may have overlooked changes in the metabolic environment of cells or high cell density. Such aspects are typically not reported or measured and may contribute to the irreproducibility of the results when repeating the assay in a different laboratory. Such irreproducibility is fueling the reproducibility debate^1,2^.

Not only do our results highlight the need for reporting experimental ‘details’ concerning culture conditions, they also show a way towards rationalizing and standardizing these. Required details would include, but not be limited to, information on the source of cell lines and passage number (or at least whether all cells used were below a certain passage number), number of cells per well at the time of seeding and throughout the assay, density/confluence throughout the assay, volume of culture media used, details of cell culture flasks used, length of assay (from time of seeding), choice of cell culture medium (including concentrations of all components) and sera (concentration used and source), and how concentrations of key nutrients (e.g. glucose and glutamine) and pH change throughout the assay. This would complement existing efforts for standardization across biomedical research^34-36^ and improve reproducibility, transparency and evaluation of the experimental data, points that are of critical concern^37^.

A number of reporting guidelines for the results of biological assays have been in existence for some time, e.g. the Minimum Information About a Microarray Experiment (MIAME) standard. MIAME is now a reporting requirement for a number of funding agencies and journals^38^. Similarly, there are now minimum reporting standards in use for metabolomics ^35^, proteomics^36^ and systems biology models^39^. Since 2008, the Minimum Information for Biological and Biomedical Investigations (MIBBI) project has acted as a repository for the many minimum reporting guidelines that have since been created; there are now over 40 for the biological and biomedical sciences (https://biosharing.org/standards/, accessed 04 July 2016).

The assay developed and discussed here, as well as the contention that it should be accompanied by metabolic analyses of the assay cell lines, should contribute to improved assay reproducibility in cell biology and drug discovery.

## 4 Acknowledgements

We thank S. E. Critchlow for guidance and B. Patel and S. Powell for providing the GLS1 inhibitor compound and sensitivity data (such as Fig. S1) for this work, as well as F. Michopoulos and H. Lewis for training, and use of the analytical instrumentation. We also thank Atilla Ting for feedback and comments on review of this manuscript. This work was supported the Biotechnology and Biological Sciences Research Council (BBSRC) CASE studentship grant in partnership with AstraZeneca (award number 1088313).

## 5 Author Contributions

MWM and HVW established the aim and strategy of the study and designed the experiments, which MWM performed. FO, RB, and CB provided advice and expertise on the design of the experiments, data acquisition and analysis. SEC, BP and SP (see acknowledgements) proposed the experimental system of glutaminase inhibition in human non-small cell lung cancer cell lines. MWM, HVW and FO wrote the paper with advice and guidance provided by RB and CB.

## 6 Competing financial interests

The authors declare that they have no competing financial interests.

